# Bullying Victimisation, Medial Occipital White Matter Development, and Psychotic-Like Experiences: A Longitudinal Diffusion MRI Study

**DOI:** 10.64898/2026.11.09.764793

**Authors:** Michael Connaughton, Orla Mitchell, Colm Healy, Aisling O’Neill, Niamh Dooley, Adi Agarwal, Ope Ogunmola, Sahar Riaz, Mary Cannon, Darren Roddy

## Abstract

Childhood bullying victimisation is robustly associated with psychotic-like experiences, but the neurodevelopmental mechanisms underlying this association remain unclear. This longitudinal study examined whether childhood bullying victimisation was associated with altered maturation of medial occipital white matter and whether bullying victimisation moderated the association between fractional anisotropy (FA) and psychotic-like experiences (PLE). A total of 96 young people (38 bullied, 58 controls) from the Adolescent Brain Development study underwent diffusion MRI at approximately 14, 16, and 20 years of age. PLE and bullying victimisation group classification were determined by expert consensus interview. FA from bilateral calcarine–cuneus, cuneus–lingual, and calcarine–lingual connections was examined using mixed-effects models. Compared with controls, the bullied group showed significantly different age-related FA trajectories in the right calcarine–cuneus connection (β = -0.40, p < .001) and the left cuneus–lingual connection (β = -0.35, p = .003). Bullying victimisation significantly moderated the association between white matter FA and PLE (β = -6.14, p = .004), indicating that the association between FA and psychotic-like experiences differed according to bullying victimisation status. This moderation effect also varied with age, as indicated by a significant bullying victimisation-by-FA-by-time interaction (β = -6.80, p = .008). Specifically, the moderating effect of bullying victimisation on the association between right calcarine–cuneus FA and PLE changed across development. These findings suggest that bullying victimisation is associated with divergent maturation of medial occipital circuitry and identify right calcarine–cuneus development as a potential marker of vulnerability to PLE following childhood bullying.

## Introduction

Bullying victimisation, characterised by repeated and targeted peer aggression, is common during adolescence and can have lasting consequences for mental health (1, 2). It is also among the most consistently identified social risk factors for psychotic-like experiences (2-5). These experiences, including hallucination-like and delusion-like phenomena, are clinically important because they are associated with increased risk for later psychotic illness, broader psychopathology, and functional impairment (6). Despite this well-established association, the neurodevelopmental mechanisms through which bullying victimisation may contribute to psychotic-like experiences remain poorly understood.

Bullying represents a particularly salient form of interpersonal adversity because it is repeated, socially threatening, and often occurs during a developmental period in which self-concept, social cognition, and threat-monitoring systems are still maturing (7). Recurrent experiences of humiliation, exclusion, intimidation, and social defeat may heighten vigilance to rejection, strengthen negative expectations about the self and others, and increase the tendency to interpret ambiguous social cues as threatening or suspicious (7). During adolescence, when neural systems remain highly responsive to environmental experience, repeated exposure to such social threat may become embedded within developing neural networks, with enduring consequences for cognition and behaviour (8-10). Identifying the neural systems through which these experiences become developmentally instantiated may therefore help explain why bullying is associated with later vulnerability to psychotic-like experiences.

Neurobiological models of adversity and psychosis have predominantly focused on stress-sensitive systems, including frontolimbic, hippocampal, and salience-network circuitry involved in stress responsivity, affect regulation, salience attribution, and cognitive control (11-14). However, psychotic-like experiences also involve disturbances in how sensory information is perceived and interpreted. Predictive-processing theories suggest that perception is shaped by both incoming sensory information and prior expectations. Psychotic-like experiences may occur when the balance between these two sources of information is disrupted, leading the brain to form inaccurate or maladaptive interpretations of sensory input (15). From this perspective, neural systems involved in the early organisation and subsequent interpretation of sensory information may represent an important, but comparatively understudied, developmental substrate linking adverse social experiences to psychosis vulnerability.

The medial occipital lobe is a compelling candidate system in which to investigate this possibility. The calcarine cortex, cuneus, and lingual gyrus form a closely situated set of visual regions spanning early and higher-order stages of visual processing (Figure 1) (16). The primary visual cortex is located predominantly along the banks of the calcarine sulcus, while the adjacent cuneus and lingual gyrus contribute to processes including visuospatial integration, motion processing, visual memory, and face-related perception (16). These functions are particularly relevant to social environments, in which facial expression, gaze, body movement, and other visual cues provide information about the intentions and behaviour of others. Altered development within this system could therefore affect the precision or stability of perceptual information before it is integrated with broader systems involved in attention, salience attribution, threat appraisal, and social meaning.

**Figure 1.**
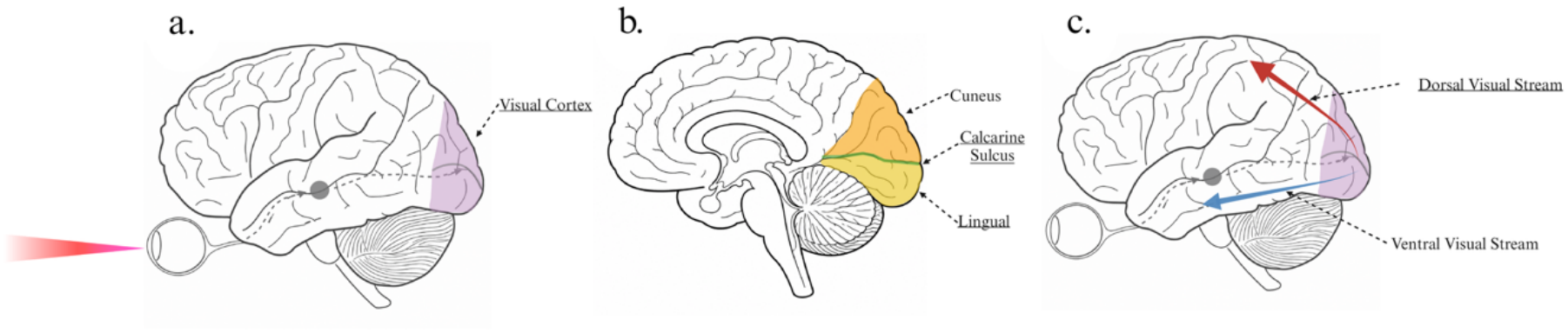
Visual pathways and the medial occipital lobe. (a) Schematic representation of the visual pathway from the eye to the occipital cortex. Visual information from the retina is transmitted via the optic nerve and retinogeniculate pathway to the lateral geniculate nucleus of the thalamus, before reaching the primary visual cortex through the optic radiations. (b) Medial view of the occipital lobe highlighting the cuneus, calcarine sulcus, and lingual gyrus. The cuneus lies superior to the calcarine sulcus and the lingual gyrus inferior to it. Primary visual cortex is located predominantly along the banks of the calcarine sulcus, while the adjacent cuneus and lingual gyrus contribute to early and higher-order visual processing, including visuospatial processing, motion processing, visual memory, and face-related perception. (c) Schematic representation of the major cortical visual processing streams extending from occipital cortex. The dorsal visual stream projects predominantly toward parietal regions and supports spatial processing, visual attention, motion processing, and visually guided action. The ventral visual stream projects predominantly toward temporal regions and supports the processing of object form, faces, and visual identity. Together, the panels situate the medial occipital lobe within the broader organisation of the visual system.

There is emerging evidence that bullying victimisation may be associated with development of these regions. Previous longitudinal structural MRI work demonstrated that greater bullying victimisation was associated with altered developmental trajectories of the cuneus and lingual gyrus across adolescence and early adulthood, reflected in age-related volumetric reductions (10). These findings suggest that the medial occipital lobe may be developmentally sensitive to interpersonal adversity. However, regional morphology represents only one aspect of neurodevelopment. Efficient visual processing also depends on coordinated communication between cortical regions, raising the possibility that bullying victimisation may be associated with altered maturation of the connections linking medial occipital regions. Characterising these developmental trajectories may therefore provide further insight into the neural mechanisms through which bullying becomes associated with later psychotic-like experiences.

Diffusion-weighted MRI provides an in vivo approach for examining the microstructural organisation of these connections (17). High Angular Resolution Diffusion Imaging (HARDI), together with constrained spherical deconvolution, improves the reconstruction of complex fibre configurations by allowing multiple fibre orientations to be modelled within a voxel (18, 19). Fractional anisotropy (FA) provides a diffusion-derived measure of white matter microstructural organisation, reflecting the degree to which water diffusion is directionally constrained within tissue (18). Across adolescence, FA generally increases with age, consistent with ongoing maturation involving changes in fibre coherence, axonal organisation, and myelination (20). Longitudinal assessment of FA therefore provides a means of testing whether bullying victimisation is associated with divergent developmental trajectories within medial occipital pathways.

The present longitudinal study examined whether development of medial occipital connectivity may contribute to the association between bullying victimisation and psychotic-like experiences across the transition from adolescence to early adulthood. Using HARDI data acquired at approximately 14, 16, and 20 years of age, we addressed two questions: first, whether bullying victimisation was associated with altered developmental trajectories of FA within bilateral calcarine–cuneus, cuneus–lingual, and calcarine–lingual connections; and second, whether medial occipital cortex FA moderated the relationship between bullying victimisation and psychotic-like experiences. Based on previous longitudinal evidence implicating medial occipital development in bullying victimisation (10, 21), it was hypothesised that bullied participants would show altered age-related FA trajectories relative to non-bullied controls. It was further hypothesised that medial occipital FA would moderate the association between bullying victimisation and psychotic-like experiences.

## Materials and Methods

### Participants

Participants were drawn from the Adolescent Brain Development (ABD) study, a longitudinal, community-based cohort recruited from mainstream primary schools in Dublin and Kildare, Ireland (22). The original cohort comprised 211 young people aged 11–13 years. During the initial phase of recruitment, approximately the first 20% of participants were enriched at a ratio of 2:1 for adolescents scoring ≥2 on the Adolescent Psychotic Symptom Screener; the remainder of the cohort was randomly sampled from the larger school-based survey population. Thus, although recruitment was community-based rather than clinical, the cohort included deliberate enrichment for psychotic-like symptoms during the early phase of recruitment.

All 211 participants in the original cohort were invited to participate in the neuroimaging component of the ABD study, which commenced 1–3 years (mean approximately 2 years) after the original clinical assessment. Participants were eligible for neuroimaging if they had no contraindications to MRI. One hundred participants completed the baseline imaging assessment. For the present study, participants contributed all usable diffusion MRI data across three imaging assessments conducted during adolescence and early adulthood; inclusion in the longitudinal analyses did not require completion of all three imaging timepoints. Participants included in the present analyses had no formal psychotic disorder or history of neurological disorder and were antipsychotic-naïve throughout the assessment period. The final analytic sample comprised 96 participants, including 38 with a reported history of childhood bullying victimisation and 58 controls.

Ethical approval was obtained from the Medical Research Ethics Committee, Beaumont Hospital, Dublin. Written parental consent and participant assent were obtained for participants younger than 18 years, with written participant consent obtained once participants reached 18 years of age.

### Clinical Measures

Participants completed diagnostic clinical interviews administered by trained raters, with recruitment and assessment procedures described previously (22). At baseline, adolescents and their parents or guardians were interviewed separately using the Schedule for Affective Disorders and Schizophrenia for School-Aged Children—Present and Lifetime Version (K-SADS-PL) (23). The K-SADS-PL was also administered at the first follow-up, although parent or guardian interviews were not conducted at follow-up assessments. At the second follow-up, participants were assessed using the Structured Clinical Interview for DSM-5 (SCID-5) (24).

*Bullying victimisation* was assessed at baseline as part of the K-SADS-PL clinical interview (23, 25). Participants and their parents or guardians were systematically asked about any history of the participant being the victim of bullying. For the present study, bullying victimisation was coded as a binary variable indicating the presence or absence of a reported history of bullying.

*Psychotic-like experiences* were assessed across all available clinical assessment waves using the psychosis sections of the K-SADS-PL and SCID-5, supplemented with items from the SOCRATES instrument (26). These assessments covered hallucination-like and delusion-like experiences. For the present analyses, PLE status was defined at the participant level using an ever/never classification across available assessment waves. Participants were classified as having experienced PLEs if one or more such experiences were identified at any assessment and as not having experienced PLEs if no such experiences were identified across available assessments. Further details of the assessment and classification procedure are provided in the Supplementary Material.

### MRI Acquisition

MRI data were acquired at the Trinity College Institute of Neuroscience, Dublin, using a Philips Intera Achieva 3.0-T MRI system with an 8-channel head coil. High-angular-resolution diffusion imaging (HARDI) was acquired using a spin-echo echo-planar imaging sequence (TE = 52 ms, TR = 11,260 ms, flip angle = 90°, field of view = 224 mm, 60 axial slices, in-plane resolution = 1.75 × 1.75 mm^2^, slice thickness = 2 mm, *b* = 1500 s/mm^2^, 61 non-collinear diffusion directions). A non-diffusion-weighted volume was also acquired. High-resolution T1-weighted anatomical images were acquired using a T1W-IR1150 sequence (TE = 3.8 ms, TR = 8.4 ms, field of view = 230 mm, in-plane resolution = 0.898 × 0.898 mm^2^, slice thickness = 0.9 mm, flip angle = 8°).

### MRI Data Processing

Diffusion MRI preprocessing and tractography were performed using ExploreDTI version 4.8.5 (27), following established processing procedures (28). Preprocessing included correction for signal drift, Gibbs ringing, echo-planar imaging distortion, and participant motion. Diffusion images were co-registered to the corresponding T1-weighted anatomical image, with appropriate rotation of the b-matrix during correction procedures (29, 30). Robust tensor estimation was performed using Robust Extraction of Kurtosis Indices with Linear Estimation (REKINDLE) (31, 32)

Whole-brain fibre reconstruction was performed using deterministic constrained spherical deconvolution tractography (18), with a 2 × 2 × 2 mm^3^ seed resolution, 1-mm step size, 60° angle threshold, and fibre lengths of 10–500 mm. Automated tractography was performed using regions of interest derived from the Automated Anatomical Labeling atlas (33). Bilateral cuneus, calcarine cortex, and lingual gyrus regions were selected to examine connections within the medial occipital lobe (Figure 2). Connections between each pair of regions were identified using the ExploreDTI “END” criterion. Bilateral calcarine–cuneus, cuneus–lingual, and calcarine–lingual connections were reconstructed, and FA was extracted from each connection for subsequent analysis.

**Figure 2.**
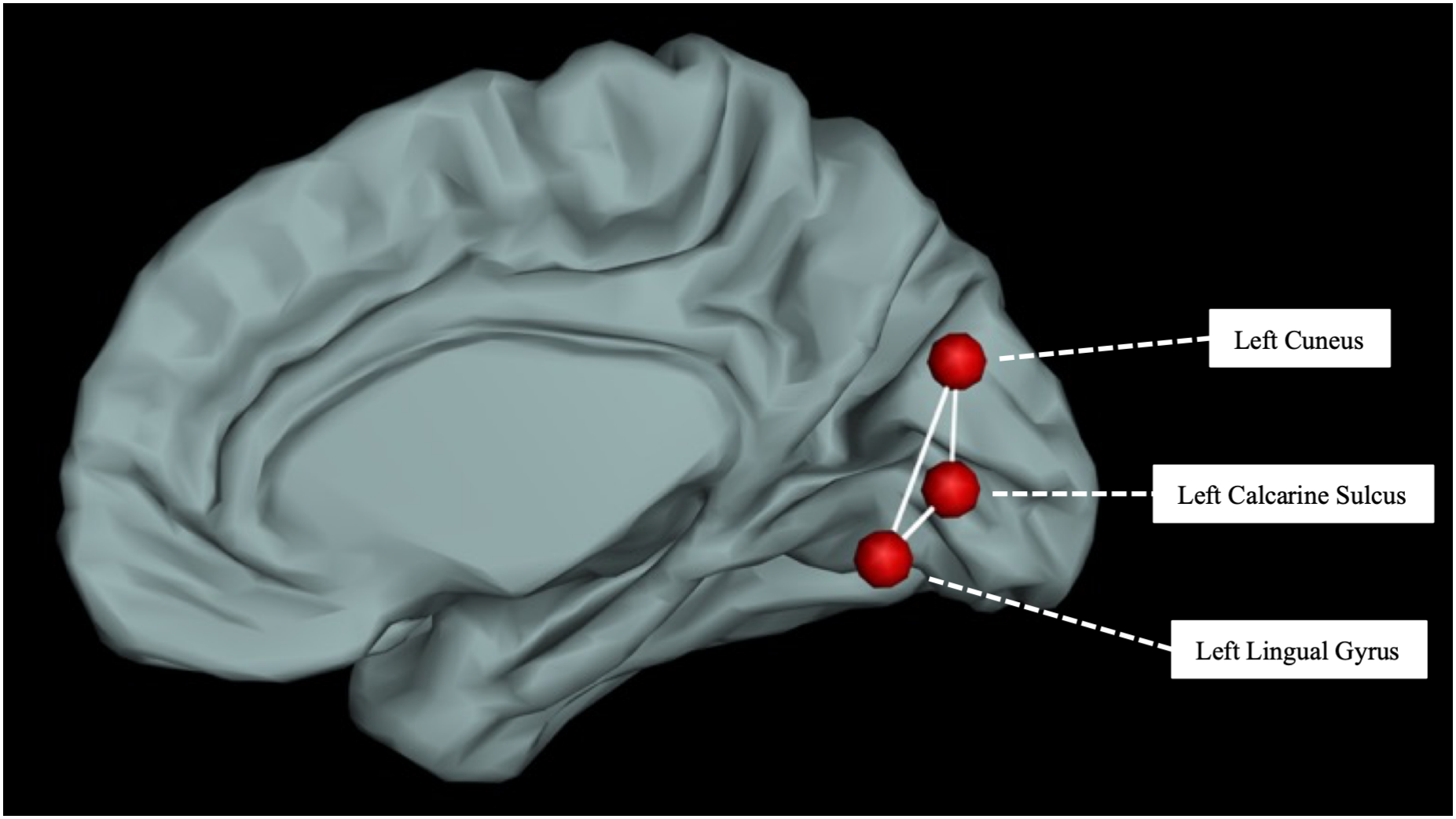
Medial occipital lobe regions of interest and inter-regional connections. Red markers indicate the selected regions of interest from the Automated Anatomical Labeling atlas: cuneus, calcarine cortex, and lingual gyrus. White lines represent the defined connections between these regions from which FA was extracted. The example shown illustrates left-hemisphere medial occipital lobe regions and their inter-regional connections.

### Statistical Analysis

For both study aims, model fitting followed a structured top-down selection procedure. Nested models were compared using likelihood ratio tests (LRTs), with more complex terms retained only when their inclusion significantly improved model fit (LRT *p* < .05) (34). Models used for comparison were fitted using maximum likelihood estimation. Final linear mixed-effects models were refitted using restricted maximum likelihood estimation for reporting of parameter estimates. A complete summary of all models tested is provided in eTable 1.

Aim 1 tested whether participants with a history of childhood bullying victimisation showed different longitudinal trajectories of medial occipital white matter FA compared with controls. FA was modelled as the longitudinal outcome using repeated observations across up to three imaging assessments. Analyses were conducted separately for the bilateral calcarine– cuneus, cuneus–lingual, and calcarine–lingual connections. Time was indexed as months since the baseline imaging assessment. For each connection, linear and quadratic representations of time were compared to determine the functional form that best characterised FA development. A quadratic time term (time^2^) was retained only when its inclusion significantly improved model fit. The primary effect of interest was the bullying victimisation-by-time interaction, which tested whether longitudinal FA trajectories differed between bullied participants and controls. The main effect of bullying victimisation was also estimated. Models included sex, total childhood stress exposure, family stress, and age at baseline as covariates. To account for multiple comparisons in Aim 1, false discovery rate correction (FDR) was applied at *q* < .05 to *p*-values associated with the bullying victimisation main effect and bullying victimisation-by-time interaction in the optimally fitted models (35, 36).

Aim 2 tested whether the association between medial occipital FA and PLEs differed according to bullying victimisation status. PLE status was modelled as the outcome using generalised linear mixed-effects models with a binomial error distribution and logit link function, implemented using the *glmer* function in *lme4* (37). Repeated FA observations across imaging assessments were incorporated within participants, with participant-level random effects accounting for within-participant dependence. The primary effect of interest was the bullying victimisation-by-FA interaction, which tested whether bullying victimisation moderated the association between FA and PLEs. A bullying victimisation-by-FA-by-time interaction was also evaluated to determine whether this moderation effect varied across development. The same top-down model-selection procedure described above was applied. FDR correction was applied at *q* < .05 to account for multiple comparisons in Aim 2 models (35, 36).

## Results

### Demographics

Full participant demographic information is summarized in Table 1. The final analytic sample included 96 participants (38 in the bullied group and 58 controls). Briefly, groups did not differ in number of available scans, age at any assessment wave, sex, socioeconomic status, family stress, total childhood stress scores or childhood DSM-5 diagnosis. Compared with controls, bullied participants were more likely to report psychotic-like experiences.

### Medial Occipital Lobe White Matter Development and Bullying Victimization

Full details of the top-down Model selection fit statistics are provided in eTable 2-4, with the final models derived from the top-down selection procedure shown in eTable 4.

Following FDR correction, significant bullying victimisation-by-time interactions remained for two medial occipital white matter connections. Compared with controls, the bullied group showed significantly different age-related FA trajectories in the right calcarine– cuneus connection (β = -0.40, 95% CI = -0.62 to -0.17, p < .001) and the left cuneus–lingual connection (β = -0.35, 95% CI = -0.57 to -0.12, p = .003). This indicates that the association between bullying victimisation and differences in developmental trajectories of medial occipital lobe white matter was present across development after accounting for age, sex, baseline age, total stress exposure, and family stress (Figure 3).

**Figure 3.**
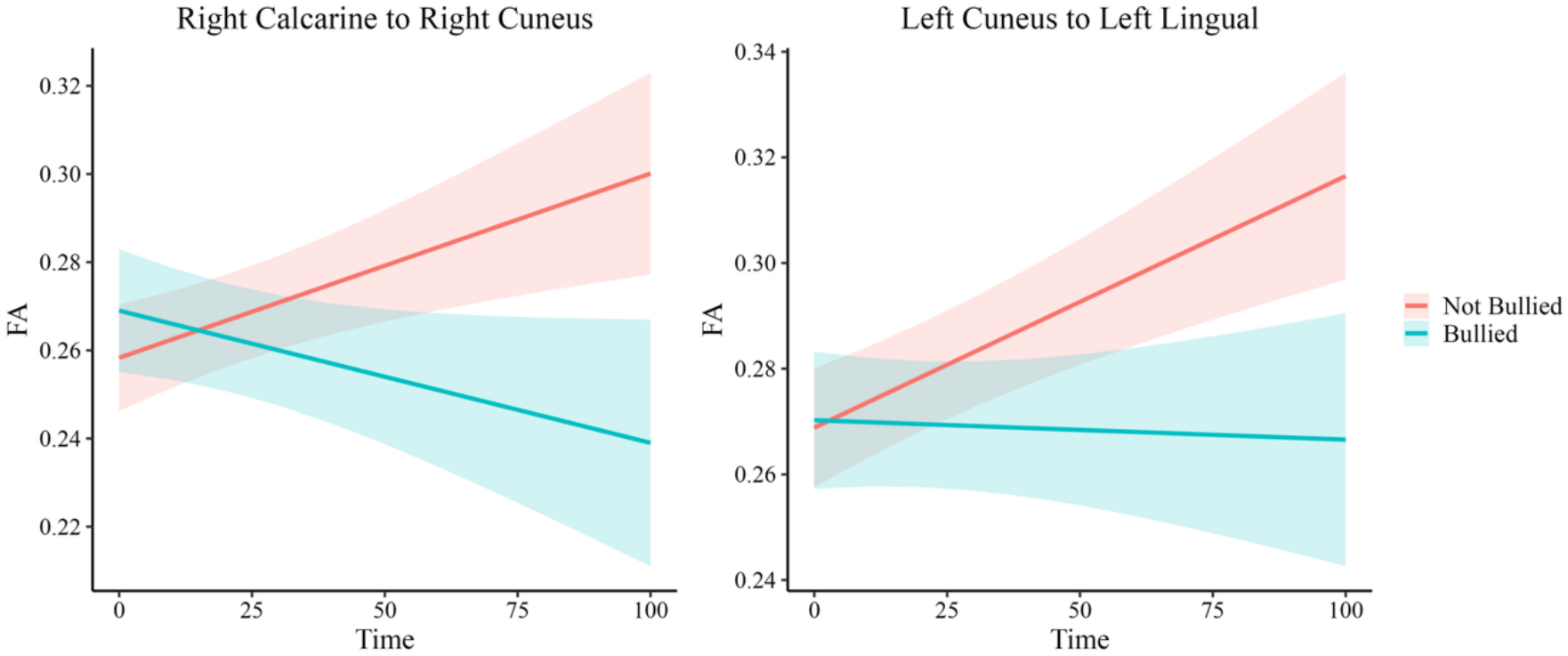
Longitudinal FA trajectories associated with bullying victimisation. Predicted fractional anisotropy (FA) trajectories for the right calcarine–right cuneus and left cuneus– left lingual white matter connections, the two tracts showing significant bullying-by-time interactions following false discovery rate (FDR) correction. Time represents months since the baseline scan. Coral lines represent participants who were not bullied, and cyan lines represent participants who were bullied. Shaded areas indicate 95% confidence intervals. Estimates are adjusted for sex, baseline age, total stress, and family stress.

There were no significant main effects of bullying victimization across these connections, suggesting that the observed associations primarily reflected differences in developmental trajectories rather than overall between-group FA differences. The full results for Study Aim 1 are reported in eTable 5.

### Bullying Victimization, Medial Occipital Lobe White Matter and Psychotic-like Experiences

FDR correction, significant effects were observed only for the right calcarine–cuneus connection. A significant bullying victimization-by-white matter interaction was observed (β = -6.14, SE = 2.13, 95% CI = -10.32 to -1.97, p = .004), indicating that the association between FA and psychotic-like experiences differed according to bullying victimization status. Importantly, this effect was further qualified by a significant bullying victimization-by-white matter-by-time interaction (β = -6.80, SE = 2.58, 95% CI = -11.86 to -1.74, p = .008), indicating that the moderating effect of bullying victimization on the association between right calcarine– cuneus FA and psychotic-like experiences varied with age (Figure 4).

**Figure 4.**
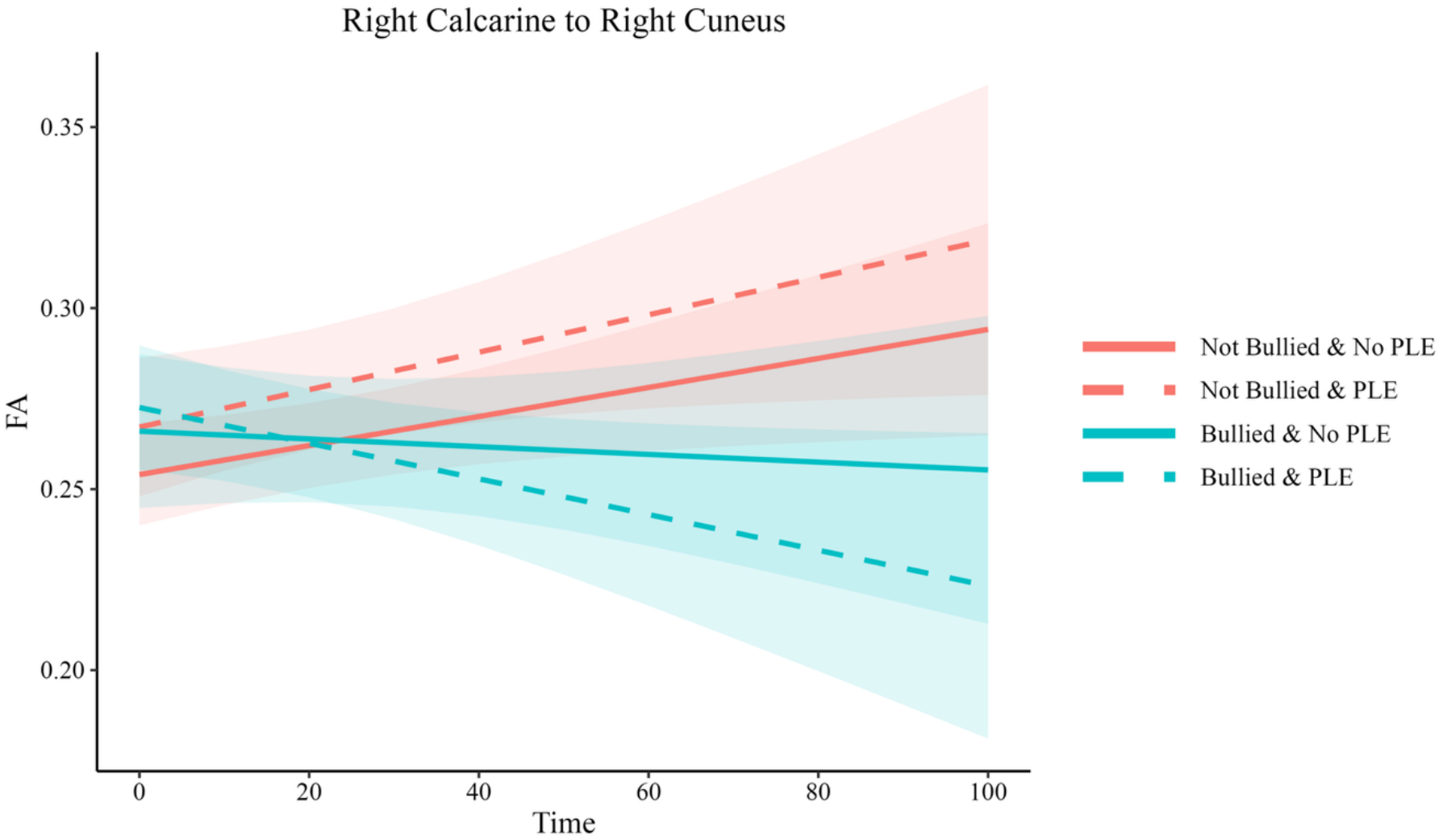
Right calcarine–cuneus FA trajectories by bullying victimisation and psychotic-like experiences. Predicted fractional anisotropy (FA) trajectories across time for the right calcarine–right cuneus white matter connection, stratified by bullying victimisation and psychotic-like experience (PLE) status. Time represents months since the baseline scan. Coral lines represent participants who were not bullied, and cyan lines represent participants who were bullied. Solid lines indicate no PLE, whereas dashed lines indicate PLE. Shaded areas indicate 95% confidence intervals. Estimates are adjusted for sex, baseline age, total stress, and family stress.

No other bullying victimization-by-white matter or bullying victimization-by-white matter-by-time interactions survived FDR correction. The full results are reported in eTable 6.

## Discussion

This three-time-point longitudinal neuroimaging study examined the relationships between bullying victimisation, psychotic-like experiences, and medial occipital lobe white matter development across adolescence and early adulthood. As hypothesised, compared to controls, the bullied group was significantly associated with, 1) altered developmental trajectories of medial occipital white matter FA, specifically right calcarine–cuneus, and left cuneus–lingual connections; and 2) right calcarine–cuneus FA moderated the association between bullying victimisation and psychotic-like experiences across adolescence and early adulthood.

These findings extend previous structural MRI work linking bullying victimisation to altered medial occipital lobe development (10), suggesting that bullying-related neurodevelopmental differences may involve not only regional morphology, but also the connectivity architecture linking visual-association regions. The following sections discuss potential mechanisms that might drive the observed associations between bullying victimisation, medial occipital white matter development, and vulnerability to psychotic-like experiences.

### Medial Occipital Circuitry as a Developmental Substrate of Bullying Victimisation

In the present study, non-bullied participants showed increasing medial occipital FA across development, whereas bullied participants showed decreasing FA over the same period. Normative diffusion studies suggest that white matter FA generally increases across adolescence, with tract-specific peaks extending into adulthood (20). This divergence may reflect altered developmental timing in this system, including accelerated maturation followed by early plateau or decline, or differences in tract composition, crossing-fibre architecture, axonal organisation, myelination, or overlapping fibre populations (38).

Taking an experience-dependent mechanistic approach, these developmental differences may relate to the perceptual demands of repeatedly monitoring socially threatening cues during adolescence. Victimised children may attend closely to facial expression, gaze direction, body movement, peer-group behaviour, and ambiguous social cues for signs of hostility, exclusion, or intimidation (39). Such experiences may place sustained demands on visual systems involved in organising and interpreting socially relevant information. The medial occipital lobe is well positioned to support these processes: the calcarine cortex contributes to early visual encoding, while the cuneus and lingual gyrus are involved in visuospatial processing, visual memory, imagery, and face-related perception (16). Through broader association pathways, visual information from these regions can be integrated with systems involved in memory, semantic processing, salience attribution, and social interpretation (16). Medial occipital circuitry may therefore be relevant to bullying victimisation because it supports the visual organisation of cues from which peer threat, exclusion, and intent are inferred.

A possible neurobiological mechanism is altered visual cortical plasticity and stabilisation. During development, visual networks remain sensitive to environmental input while progressively consolidating into more efficient, specialised, and structurally organised circuits (40, 41). This transition is supported by coordinated processes including maturation of inhibitory signalling, refinement of local cortical circuits, myelin-related signalling, and excitation–inhibition balance (40, 41). Early-life stress has been associated with alterations in cortical plasticity, network refinement, and white matter organisation (42, 43), which may prolong or shift sensitive periods of visual cortical development and have lasting consequences for visual network structure and function (44). Although preliminary, the present findings raise the possibility that interpersonal stress during neurodevelopmentally sensitive periods may be associated with altered maturation of medial occipital pathways involved in visual-association processing.

### Medial Occipital Connectivity, Predictive Processing, and Psychotic-like Experiences

The moderation finding provides a potential link between altered medial occipital development and psychotic-like experiences. Bullying victimisation was more strongly associated with psychotic-like experiences among individuals with lower FA in the right calcarine–cuneus connection, suggesting that variation within this pathway may be relevant to psychosis vulnerability. The anatomical location of this finding is notable. The medial occipital lobe is a highly interconnected system supporting both early and higher-order visual processing, with local connections enabling communication between the cuneus, lingual gyrus, and adjacent visual cortex (16, 45, 46). As illustrated in Figure 1, visual input reaching calcarine cortex is progressively organised across medial occipital and visual-association regions before contributing to broader perceptual processing. Altered maturation of calcarine– cuneus, cuneus–lingual, and calcarine–lingual connections could therefore affect the fidelity, timing, or coordination of visual information passing through this system. Importantly, disturbances of early visual processing are well documented in schizophrenia, and diffusion MRI studies have linked reduced integrity of visual-system white matter, particularly the optic radiations, with impaired early visual responses (47, 48). More recent work has also identified visual-processing abnormalities in individuals at clinical high risk and during first-episode psychosis, suggesting that disruption of visual information processing is not restricted to chronic illness (49, 50).

These findings are relevant to predictive-processing models, in which perception depends on recurrent comparisons between incoming sensory information and predictions generated at higher levels of the perceptual hierarchy (15, 51). A mismatch between predicted and observed input generates a prediction error, which is used to update the brain’s model of the environment; the impact of that error depends on the precision, or reliability, attributed to the underlying sensory evidence (15, 51). Altered maturation of medial occipital connections could therefore be relevant on the sensory side of this computation. If visual information is transmitted or integrated less consistently between early and association regions, the sensory representation against which predictions are evaluated may be less stable or precise. Prediction errors derived from that information could consequently be generated, propagated, or weighted atypically, altering the extent to which sensory evidence can appropriately update perceptual interpretations. There is direct evidence that this process is disrupted in psychosis: reduced visual mismatch responses, widely interpreted as indices of visual prediction error, have been demonstrated in schizophrenia using both simple visual features and more complex stimuli (52, 53), and a recent meta-analysis of ten studies found a moderate reduction in visual mismatch negativity in schizophrenia (54). Thus, although FA does not measure prediction-error signalling directly, the present findings identify developmental differences within circuitry positioned to influence the visual evidence upon which such signalling depends.

Bullying may then influence what happens when this visual evidence is interpreted. Bullying victimisation can strengthen expectations of hostility and has been associated with paranoid thinking and overestimation of threat in neutral social situations (39, 55). In this context, altered medial occipital processing and bullying-related expectations may act together. If ambiguous visual information provides a weaker or less reliable constraint on perception, pre-existing expectations of interpersonal threat may exert greater influence over how facial expressions, gaze, body movements, or peer interactions are interpreted. Conversely, atypically weighted prediction errors could cause otherwise innocuous sensory events to acquire excessive significance and promote maladaptive belief updating, as proposed in computational accounts of psychosis (15, 51). This provides a possible interpretation of the right calcarine– cuneus moderation effect: developmental variation in a pathway linking early and associative visual processing may alter the sensory evidence available for perceptual inference, while bullying shapes the expectations against which that evidence is evaluated. An imbalance between these processes could increase the likelihood that ambiguous social information is experienced as hostile, unusually salient, or self-referential, contributing to suspiciousness, paranoid ideation, and other psychotic-like experiences. This interpretation remains provisional because visual function, perceptual precision, and prediction-error signalling were not directly measured in the present study.

### Strengths and Limitations

The strengths of this study are several. First, participants were recruited from the general population and were in full-time education, antipsychotic-naïve, non-smoking, and free from neurological disorder. This reduces the likelihood that the observed associations were driven by such confounders (56, 57). Second, psychotic-like experiences were assessed using detailed clinical interviews, with interview-derived symptoms verified through expert consensus ratings. This is a strength because interview-based assessment can improve the validity of psychotic-experience classification relative to screening instruments alone (58). Third, the use of constrained spherical deconvolution-based tractography allowed improved modelling of complex fibre architecture relative to diffusion tensor imaging approaches (59), which is important when reconstructing complex white matter fibres, such as those within the medial occipital lobe.

Several limitations should temper interpretation. The sample size was modest for longitudinal neuroimaging, increasing uncertainty around effect estimates and requiring replication (60). Bullying victimization was treated as a dichotomous exposure, which may obscure biologically important variation in severity, chronicity, developmental timing, relational context, and subjective distress (61). In addition, FA is a biologically non-specific measure and may reflect multiple underlying tissue properties, including fibre coherence, myelination, axonal organisation, and crossing-fibre architecture (62). As such, these findings should be interpreted as evidence of bullying-related differences in diffusion-derived white matter development, rather than direct evidence of a single underlying cellular mechanism. A further limitation is that psychotic-like experiences were analysed as a broad phenotype (63), meaning that the present study could not determine whether bullying-related neurodevelopment is differentially associated with specific symptom dimensions, such as hallucination-like experiences, suspiciousness, or threat-related interpretations.

## Conclusion

This longitudinal diffusion MRI study provides evidence that bullying victimisation is associated with divergent development of medial occipital white matter across adolescence and early adulthood. Altered FA trajectories were observed across multiple connections linking the calcarine cortex, cuneus, and lingual gyrus, indicating that bullying-related neurodevelopmental differences extend to circuitry supporting early and higher-order visual processing. Importantly, lower FA within the right calcarine–cuneus connection strengthened the association between bullying victimisation and psychotic-like experiences, identifying this pathway as a potential marker of vulnerability among bullied young people. Together, these findings implicate medial occipital circuitry as a previously under-recognised neurodevelopmental substrate linking interpersonal adversity with psychosis vulnerability and raise the possibility that altered development of visual-processing systems may influence how perceptual information is organised and interpreted following bullying.

## CRediT authorship contribution statement

Michael Connaughton: Conceptualization, Methodology, Formal analysis, Investigation, Data curation, Visualization, Writing - original draft, Writing - review … editing, Project administration; Orla Mitchell: Writing - review … editing; Colm Healy: Investigation; Aisling O’Neill: Investigation; Niamh Dooley: Investigation; Adi Agarwal: Visualization, Writing - review … editing; Ope Ogunmola: dMRI image QC, Writing - review … editing; Sahar Riaz: Writing - review … editing; Mary Cannon: Investigation; Darren Roddy: Funding acquisition, Writing - review … editing.

