## Supplemental Material for "Bullying Victimisation, Medial Occipital White Matter Development, and Psychotic-Like Experiences: A Longitudinal Diffusion MRI Study"

*Connaughton et al.,*

### 1. Methods

#### 1.1. Stress Exposure and Family Stress Variables

Total childhood stress exposure was calculated from the K-SADS-PL as the cumulative number of endorsed stressors reported during the baseline diagnostic interview, excluding bullying Victimisation. Stressors included parental separation after birth; hospital admission of one night or more; surgical or invasive medical procedures; childhood physical or sexual abuse; maternal, paternal, or sibling diagnosed mental disorder; involvement in a car accident, excluding minor incidents; other accidents; experience of a fire; witnessing a natural disaster; witnessing or being the victim of a violent crime; witnessing domestic violence in the home; death of a parent, sibling, grandparent, cousin, aunt, uncle, friend, teacher, close family friend, or known parent of a friend; and any reported experience in foster or institutional care.

Family stress was coded separately from K-SADS-PL interview data to capture stressors occurring within the family environment. This variable included parental separation, maternal mental illness, paternal mental illness, sibling mental illness, parental death, sibling death, domestic violence in the home, and foster or institutional care.

#### 1.2. Classification Criteria for Psychotic-like Experiences for the ABD Study

The psychosis subsection of the K-SADS, together with additional SOCRATES items, was used to assess hallucination-like experiences and delusional thinking. Responses describing possible psychotic phenomena were classified as Definite Psychotic-like Experience, Weak Psychotic-like Experience, or Not a Psychotic-like Experience. For the present study, participants were classified as having psychotic-like experiences if at least one reported experience was rated by consensus as either a Definite Psychotic-like Experience or a Weak Psychotic-like Experience. Examples of the criteria used to distinguish definite, weak, and non-psychotic-like experiences are provided in the table below.

| Domain | Definite Psychotic-like Experience | Weak Psychotic-like Experience | Not Classified as a Psychotic-like Experience |
|---|---|---|---|
| **Hallucinations** | | | |
| Auditory verbal phenomena | Hearing one or more voices saying at least one word. Hearing whispering or indistinct voices at normal volume or shouting. Hearing one's name being called, but only when attributed to a delusional belief about another entity calling the name, such as a ghost. Brief episodes of hearing one's own thoughts aloud when associated with delusional ideation and experienced as distressing or disorganising. | Brief episodes of hearing one's own thoughts aloud when not associated with delusional ideation and neither distressing nor disorganising. | Hypnagogic and hypnopompic auditory verbal experiences. |
| Non-verbal auditory phenomena | Hearing non-verbal sounds that are experienced as distressing or disorganising. Hearing non-verbal sounds associated with delusional ideation. | Brief auditory experiences such as hearing music or other non-verbal sounds, including footsteps or knocking sounds, in the absence of associated delusional ideation. | Hypnagogic and hypnopompic non-verbal auditory experiences. |
| Non-auditory perceptual phenomena | Seeing figures or forms that are not there, such as ghosts, human forms, aliens, or the devil. Tactile sensations associated with delusional attribution. | Recurrent experiences of smelling food that is not there. | Hypnagogic and hypnopompic visual experiences. Visual illusions. Occasional experiences of smells or tastes without associated distress. |
| **Delusions** | | | |
| Unusual thoughts and beliefs | Delusional beliefs pertaining to any hallucinatory experiences in Domains 1–3. Definite and fixed beliefs about being watched by a person, entity, or organisation. Recurrent and unfounded paranoid ideas that other people are criticising the individual. Beliefs that non-human entities, such as ghosts, spirits, aliens, or the devil, are communicating directly with the individual. Unshakable nihilistic beliefs. Mind reading, by self or others, when accompanied by paranoid beliefs that the individual has been singled out to have their mind read for a negative or nefarious purpose. | Vague sense or thought that the individual is being watched. Mind reading, by self or others, when not accompanied by a belief that the individual has been singled out to have their mind read for a negative or nefarious purpose. Magical thinking, such as a belief that the individual can predict the future, when not accompanied by distress or leading to disorganisation. | Thoughts and beliefs about being watched, judged, or criticised by others that occur due to self-consciousness. Beliefs that ghosts, spirits, or aliens are responsible for or can influence experiences when such beliefs are aligned with normative cultural or subcultural beliefs. Subcultural beliefs that the world is coming to an end. Subcultural conspiracy beliefs. |

#### 1.3. Tested Models for Project Statistical Analyses

**eTable 1.** Candidate Mixed-Effects Models for Bullying-Related Medial Occipital White Matter Development and Psychotic-Like Experiences.

**Study Aim 1:** Medial Occipital Visual-Association Circuit White Matter Development in Children Exposed to Bullying Victimisation

| | |
|---|---|
| Aim 1: Linear Model | FA ~ $b_1$(group) + $b_2$(group*time) + $b_3$(time) + $b_4$(sex) + $b_5$(total childhood stress) + $b_6$(family stress) + $b_7$(age at baseline) + (1\|subject) |
| Aim 1: Quadratic Model | FA ~ $b_1$(group) + $b_2$(group*time) + $b_3$(time) + $b_4$(time$^2$) + $b_5$(sex) + $b_6$(total childhood stress) + $b_7$(family stress) + $b_8$(age at baseline) + (1\|subject) |
| Aim 1: Linear simple Model | FA ~ $b_1$(group) + $b_2$(time) + $b_3$(sex) + $b_4$(total childhood stress) + $b_5$(family stress) + $b_6$(age at baseline) + (1\|subject) |
| Aim 1: Quadratic simple Model | FA ~ $b_1$(group) + $b_2$(time) + $b_3$(time$^2$) + $b_4$(sex) + $b_5$(total childhood stress) + $b_6$(family stress) + $b_7$(age at baseline) + (1\|subject) |
| Aim 1: Linear complex Model | FA ~ $b_1$(group) + $b_2$(group*time) + $b_3$(time) + $b_4$(sex) + $b_5$(total childhood stress) + $b_6$(family stress) + $b_7$(age at baseline) + (1\|subject) |
| Aim 1: Quadratic complex Model | FA ~ $b_1$(group) + $b_2$(group*time) + $b_3$(time) + $b_4$(time$^2$) + $b_5$(sex) + $b_6$(total childhood stress) + $b_7$(family stress) + $b_8$(age at baseline) + (1\|subject) |

**Study Aim 2:** Bullying Victimisation, Medial Occipital Visual-Association Circuit FA and Psychotic-like experiences

| | |
|---|---|
| Aim 2: Linear simple Model | PE ~ $b_1$(group* WM FA) + $b_2$(time) + $b_3$(sex) + $b_4$(total childhood stress) + $b_5$(family stress) + $b_6$(age at baseline) + (1\|subject) |
| Aim 2: Quadratic simple Model | PE ~ $b_1$(group* WM FA) + $b_2$(time) + $b_3$(time$^2$) + $b_4$(sex) + $b_5$(total childhood stress) + $b_6$(family stress) + $b_7$(age at baseline) + (1\|subject) |
| Aim 2: Linear complex Model | PE ~ $b_1$(group*time* WM FA) + $b_2$(time) + $b_3$(sex) + $b_4$(total childhood stress) + $b_5$(family stress) + $b_6$(age at baseline) + (1\|subject) |
| Aim 2: Quadratic complex Model | PE ~ $b_1$(group*time* WM FA) + $b_2$(time) + $b_3$(time$^2$) + $b_4$(sex) + $b_5$(total childhood stress) + $b_6$(family stress) + $b_7$(age at baseline) + (1\|subject) |

**Legend:** PE = psychotic-like experience, WM = white matter, FA = fractional anisotropy, (1|subject) = individual random effects of intercept.

### 2. Results

#### 2.1. Model Fit Statistics

**eTable 2: Fit indices of Linear vs Quadratic Models.**

| | AIC | | LL | | LRT (p-values) |
|---|---|---|---|---|---|
| **DV** | **Linear Model** | **Quadratic Model** | **Linear Model** | **Quadratic Model** | **Linear vs Quadratic Models** |
| **Aim 1** | | | | | |
| Left Calcarine to Left Cuneus | 553.60 | 554.71 | -266.80 | -266.36 | 0.8889 |
| Right Calcarine to Right Cuneus | 500.70 | 502.68 | -240.35 | -240.34 | 0.8818 |
| Left Cuneus to Left Lingual | 591.63 | 596.94 | -269.08 | -269.06 | 0.8432 |
| Right Cuneus to Right Lingual | 579.00 | 580.58 | -279.50 | -279.29 | 0.5129 |
| Left Calcarine to Left Lingual | 579.35 | 578.27 | -279.68 | -278.13 | 0.07912 |
| Right Calcarine to Right Lingual | 589.17 | 590.42 | -284.58 | -284.21 | 0.3856 |
| **Aim 2** | | | | | |
| Left Calcarine to Left Cuneus | 229.88 | 225.77 | -101.939 | -98.884 | 0.01344 |
| Right Calcarine to Right Cuneus | 215.58 | 215.31 | -94.789 | -93.655 | 0.132 |
| Left Cuneus to Left Lingual | 232.37 | 232.90 | -106.19 | -105.45 | 0.2248 |
| Right Cuneus to Right Lingual | 243.13 | 235.91 | -108.56 | -103.96 | 0.002403 |
| Left Calcarine to Left Lingual | 238.40 | 229.43 | -106.20 | -100.72 | 0.0009258 |
| Right Calcarine to Right Lingual | 244.49 | 244.43 | -109.25 | -108.22 | 0.1515 |

**Legend:** DV = dependent variable, AIC = Akaike Information Criterion, LL = Log-Likelihood, LRT = Likelihood Ratio Test.

**eTable 3: Fit indices of Fixed Effects Models.**

| | AIC | | LL | | LRT (p-values) |
|---|---|---|---|---|---|
| **DV** | **Simple Model** | **Complex Model** | **Simple Model** | **Complex Model** | **Simple Model vs. Complex Model** |
| **Aim 1** | | | | | |
| Left Calcarine to Left Cuneus | 555.6 | 553.6 | -268.8 | -266.8 | 0.04567 |
| Right Calcarine to Right Cuneus | 511.01 | 500.70 | -246.50 | -240.35 | 0.0004504 |
| Left Cuneus to Left Lingual | 565.28 | 558.16 | -273.64 | -269.08 | 0.002529 |
| Right Cuneus to Right Lingual | 581.11 | 579.00 | -281.56 | -279.50 | 0.04267 |
| Left Calcarine to Left Lingual | 578.14 | 579.35 | -280.07 | -279.68 | 0.3739 |
| Right Calcarine to Right Lingual | 588.42 | 589.17 | -285.21 | -284.58 | 0.2633 |
| **Aim 2** | | | | | |
| Left Calcarine to Left Cuneus | 224.87 | 225.77 | -101.436 | -98.884 | 0.1643 |
| Right Calcarine to Right Cuneus | 219.92 | 215.58 | -100.961 | -94.789 | 0.01497 |
| Left Cuneus to Left Lingual | 232.37 | 229.19 | -106.19 | -101.59 | 0.02692 |
| Right Cuneus to Right Lingual | 232.31 | 235.91 | -105.15 | -103.96 | 0.4949 |
| Left Calcarine to Left Lingual | 225.78 | 229.43 | -101.89 | -100.72 | 0.5036 |
| Right Calcarine to Right Lingual | 238.95 | 244.43 | -108.48 | -108.22 | 0.9146 |

**Legend:** DV = dependent variable, AIC = Akaike Information Criterion, LL = Log-Likelihood, LRT = Likelihood Ratio Test.

**eTable 4: Results of Top-Down Mixed Model Selection: Optimal Models**

| DV | Linear Simple Model | Linear Complex Model | Quadratic Simple Model | Quadratic Complex Model |
|---|---|---|---|---|
| **Aim 1** | | | | |
| Left Calcarine to Left Cuneus | | ☑ | | |
| Right Calcarine to Right Cuneus | | ☑ | | |
| Left Cuneus to Left Lingual | | ☑ | | |
| Right Cuneus to Right Lingual | | ☑ | | |
| Left Calcarine to Left Lingual | ☑ | | | |
| Right Calcarine to Right Lingual | ☑ | | | |
| **Aim 2** | | | | |
| Left Calcarine to Left Cuneus | | | ☑ | |
| Right Calcarine to Right Cuneus | | ☑ | | |
| Left Cuneus to Left Lingual | | ☑ | | |
| Right Cuneus to Right Lingual | | | ☑ | |
| Left Calcarine to Left Lingual | | | ☑ | |
| Right Calcarine to Right Lingual | ☑ | | | |

#### 2.2. Results of Optimal Models

**eTable 5: Results of Optimal Model (Study Aim 1)**

| ROI | Intercept | | | | Time | | | | Age at Baseline | | | | Sex | | | | Total Stress | | | | Family Stress | | | | Bully Victim Group | | | | Bully Victim Group*Time | | | | |
|---|---|---|---|---|---|---|---|---|---|---|---|---|---|---|---|---|---|---|---|---|---|---|---|---|---|---|---|---|---|---|---|---|---|
| | β | SE | *t* | *p* | β | SE | *t* | *p* | β | SE | *t* | *p* | β | SE | *t* | *p* | β | SE | *t* | *p* | β | SE | *t* | *p* | β | SE | *t* | *p* | β | SE | *t* | *p* | *fdr p* |
| Left Calcarine to Left Cuneus | -0.20590 | 0.14669 | -1.404 | 0.1626 | 0.16509 | 0.07488 | 2.205 | 0.0290 | 0.13499 | 0.07600 | 1.776 | 0.0789 | 0.31110 | 0.14966 | 2.079 | 0.0404 | 0.03160 | 0.04443 | 0.711 | 0.4779 | -0.05636 | 0.14008 | -0.402 | 0.6879 | -0.14020 | 0.15243 | -0.920 | 0.3600 | -0.22927 | 0.11657 | -1.967 | 0.0510 | 0.0510 |
| Right Calcarine to Right Cuneus | -0.11145 | 0.12072 | -0.923 | 0.357604 | 0.26152 | 0.07208 | 3.628 | 0.000383 | 0.06697 | 0.05923 | 1.131 | 0.261259 | 0.20234 | 0.11578 | 1.748 | 0.084232 | 0.01839 | 0.04183 | 0.440 | 0.660657 | -0.09999 | 0.13060 | -0.766 | 0.444845 | -0.25901 | 0.11952 | -2.167 | 0.032829 | -0.39801 | 0.11345 | -3.508 | 0.000579 | 0.0034 |
| Left Cuneus to Left Lingual | -0.12346 | 0.14904 | -0.828 | 0.40884 | 0.30823 | 0.07337 | 4.201 | 0.000046 | 0.13992 | 0.07825 | 1.788 | 0.07690 | 0.25026 | 0.15347 | 1.631 | 0.10633 | 0.03785 | 0.04434 | 0.854 | 0.39441 | -0.07347 | 0.13981 | -0.526 | 0.59987 | -0.30406 | 0.15617 | -1.947 | 0.05442 | -0.34672 | 0.11489 | -3.018 | 0.00299 | 0.0089 |
| Right Cuneus to Right Lingual | 0.08492 | 0.15286 | 0.556 | 0.5794 | 0.09756 | 0.07837 | 1.245 | 0.2151 | 0.17131 | 0.07900 | 2.168 | 0.0326 | 0.21337 | 0.15489 | 1.378 | 0.1716 | -0.07142 | 0.04663 | -1.531 | 0.1274 | -0.17692 | 0.14597 | -1.212 | 0.2271 | -0.01177 | 0.15761 | -0.075 | 0.9406 | -0.24583 | 0.12230 | -2.010 | 0.0462 | 0.3870 |
| Left Calcarine to Left Lingual | -0.04392 | 0.15984 | -0.275 | 0.7839 | 0.04867 | 0.05747 | 0.847 | 0.3985 | 0.06293 | 0.08535 | 0.737 | 0.4628 | 0.25117 | 0.16782 | 1.497 | 0.1380 | -0.00781 | 0.04526 | -0.172 | 0.8633 | 0.03345 | 0.14276 | 0.234 | 0.8150 | -0.31057 | 0.16997 | -1.827 | 0.0708 | N/A | N/A | N/A | N/A | N/A |
| Right Calcarine to Right Lingual | 0.20009 | 0.15497 | 1.291 | 0.1988 | 0.06324 | 0.06193 | 1.021 | 0.3087 | 0.11857 | 0.07954 | 1.491 | 0.1395 | -0.02301 | 0.15577 | -0.148 | 0.8829 | -0.08459 | 0.04795 | -1.764 | 0.0794 | 0.03669 | 0.15071 | 0.243 | 0.8079 | -0.07075 | 0.15877 | -0.446 | 0.6569 | N/A | N/A | N/A | N/A | N/A |

**Legend**: ROI = region of interest, β = Estimate, SE = Standard error, *t* = t value, *p* = p value, *fdr p* = fdr correct p value.

**eTable 6: Results of Optimal Model (Study Aim 2)**

| ROI | Intercept | | | | Time | | | | Time² | | | | Age at Baseline | | | | Sex | | | | Total Stress | | | | Family Stress | | | |
|---|---|---|---|---|---|---|---|---|---|---|---|---|---|---|---|---|---|---|---|---|---|---|---|---|---|---|---|---|
| | β | SE | z | P | β | SE | z | p | β | SE | z | p | β | SE | z | p | β | SE | z | p | β | SE | z | p | β | SE | z | p |
| Left Calcarine to Left Cuneus | -12.517 | 3.998 | -3.131 | 0.001 | -2.60610 | 1.170 | -2.226 | 0.026 | 1.798 | 0.944 | 1.904 | 0.056 | -0.848 | 1.123 | -0.755 | 0.450 | 2.199 | 2.274 | 0.967 | 0.333 | -0.610 | 0.426 | -1.431 | 0.152 | 2.645 | 1.751 | 1.510 | 0.131 |
| Right Calcarine to Right Cuneus | -13.250 | 3.444 | -3.847 | 0.001 | -3.38617 | 1.473 | -2.298 | 0.021 | N/A | N/A | N/A | N/A | -0.981 | 1.061 | -0.925 | 0.354 | 1.195 | 1.935 | 0.618 | 0.536 | -0.083 | 0.385 | -0.216 | 0.828 | 1.783 | 1.456 | 1.225 | 0.220 |
| Left Cuneus to Left Lingual | -10.219 | 2.546 | -4.013 | 0.001 | -3.02020 | 1.192 | -2.533 | 0.011 | N/A | N/A | N/A | N/A | -0.417 | 0.824 | -0.506 | 0.612 | 1.426 | 1.654 | 0.862 | 0.388 | -0.667 | 0.390 | -1.711 | 0.087 | 2.223 | 1.328 | 1.674 | 0.094 |
| Right Cuneus to Right Lingual | -12.813 | 3.853 | -3.325 | 0.001 | -2.30530 | 1.053 | -2.187 | 0.028 | 1.908 | 0.961 | 1.984 | 0.047 | -1.083 | 1.193 | -0.908 | 0.363 | 1.548 | 2.061 | 0.751 | 0.452 | 0.133 | 0.347 | 0.385 | 0.700 | 1.176 | 1.236 | 0.951 | 0.341 |
| Left Calcarine to Left Lingual | -15.245 | 5.088 | -2.996 | 0.002 | -2.94920 | 1.299 | -2.269 | 0.023 | 2.592 | 1.313 | 1.973 | 0.048 | -0.873 | 1.133 | -0.770 | 0.441 | 2.644 | 2.509 | 1.054 | 0.291 | -0.607 | 0.480 | -1.263 | 0.206 | 3.262 | 2.142 | 1.523 | 0.127 |
| Right Calcarine to Right Lingual | -4.010 | 1.969 | -2.036 | 0.041 | -0.82248 | 0.457 | -1.798 | 0.072 | 0.535 | 0.385 | 1.388 | 0.165 | -0.512 | 0.587 | -0.873 | 0.382 | 1.388 | 1.132 | 1.226 | 0.220 | -0.089 | 0.220 | -0.404 | 0.686 | 0.220 | 0.747 | 0.295 | 0.768 |

| ROI | Bully Victim Group | | | | White Matter Tract | | | | Bully Victim Group × WM | | | | | Bully Victim Group × Time | | | | WM × Time | | | | Bully Victim Group × WM × Time | | | | |
|---|---|---|---|---|---|---|---|---|---|---|---|---|---|---|---|---|---|---|---|---|---|---|---|---|---|---|
| | β | SE | z | p | β | SE | z | p | β | SE | z | p | fdr p | β | SE | z | p | β | SE | z | p | β | SE | z | p | fdr p |
| Left Calcarine to Left Cuneus | 18.414 | 5.655 | 3.256 | 0.001 | 3.125 | 1.452 | 2.151 | 0.031 | -4.627 | 2.064 | -2.241 | 0.025 | 0.063 | N/A | N/A | N/A | N/A | N/A | N/A | N/A | N/A | N/A | N/A | N/A | N/A | N/A |
| Right Calcarine to Right Cuneus | 19.948 | 4.638 | 4.301 | <0.0001 | 3.151 | 1.015 | 3.103 | 0.001 | -6.144 | 2.132 | -2.882 | 0.003 | 0.018 | 1.775 | 1.811 | 0.980 | 0.327 | 3.031 | 1.175 | 2.578 | 0.009 | -6.800 | 2.581 | -2.634 | 0.008 | 0.048 |
| Left Cuneus to Left Lingual | 18.068 | 3.790 | 4.766 | <0.001 | 2.087 | 0.846 | 2.466 | 0.013 | -2.569 | 1.267 | -2.028 | 0.042 | 0.063 | 2.299 | 1.366 | 1.682 | 0.092 | 2.172 | 0.949 | 2.288 | 0.022 | -2.183 | 1.336 | -1.634 | 0.102 | 0.204 |
| Right Cuneus to Right Lingual | 17.375 | 4.789 | 3.628 | 0.000 | 1.995 | 1.028 | 1.941 | 0.052 | -0.620 | 1.268 | -0.489 | 0.624 | 0.624 | N/A | N/A | N/A | N/A | N/A | N/A | N/A | N/A | N/A | N/A | N/A | N/A | N/A |
| Left Calcarine to Left Lingual | 20.655 | 6.258 | 3.300 | 0.001 | 2.801 | 1.292 | 2.167 | 0.030 | -4.228 | 1.998 | -2.116 | 0.034 | 0.063 | N/A | N/A | N/A | N/A | N/A | N/A | N/A | N/A | N/A | N/A | N/A | N/A | N/A |
| Right Calcarine to Right Lingual | 4.223 | 2.382 | 1.773 | 0.076 | 0.621 | 0.499 | 1.243 | 0.213 | -1.047 | 0.737 | -1.420 | 0.155 | 0.186 | N/A | N/A | N/A | N/A | N/A | N/A | N/A | N/A | N/A | N/A | N/A | N/A | N/A |

**Legend**: ROI = region of interest, β = Estimate, SE = Standard error, *z* = z value, *p* = p value, *fdr p* = fdr correct p value.

#### 2.3. Figures of Optimal Models

**eFigure 1:** Medial Occipital White Matter FA Trajectories by Bullying Victimisation

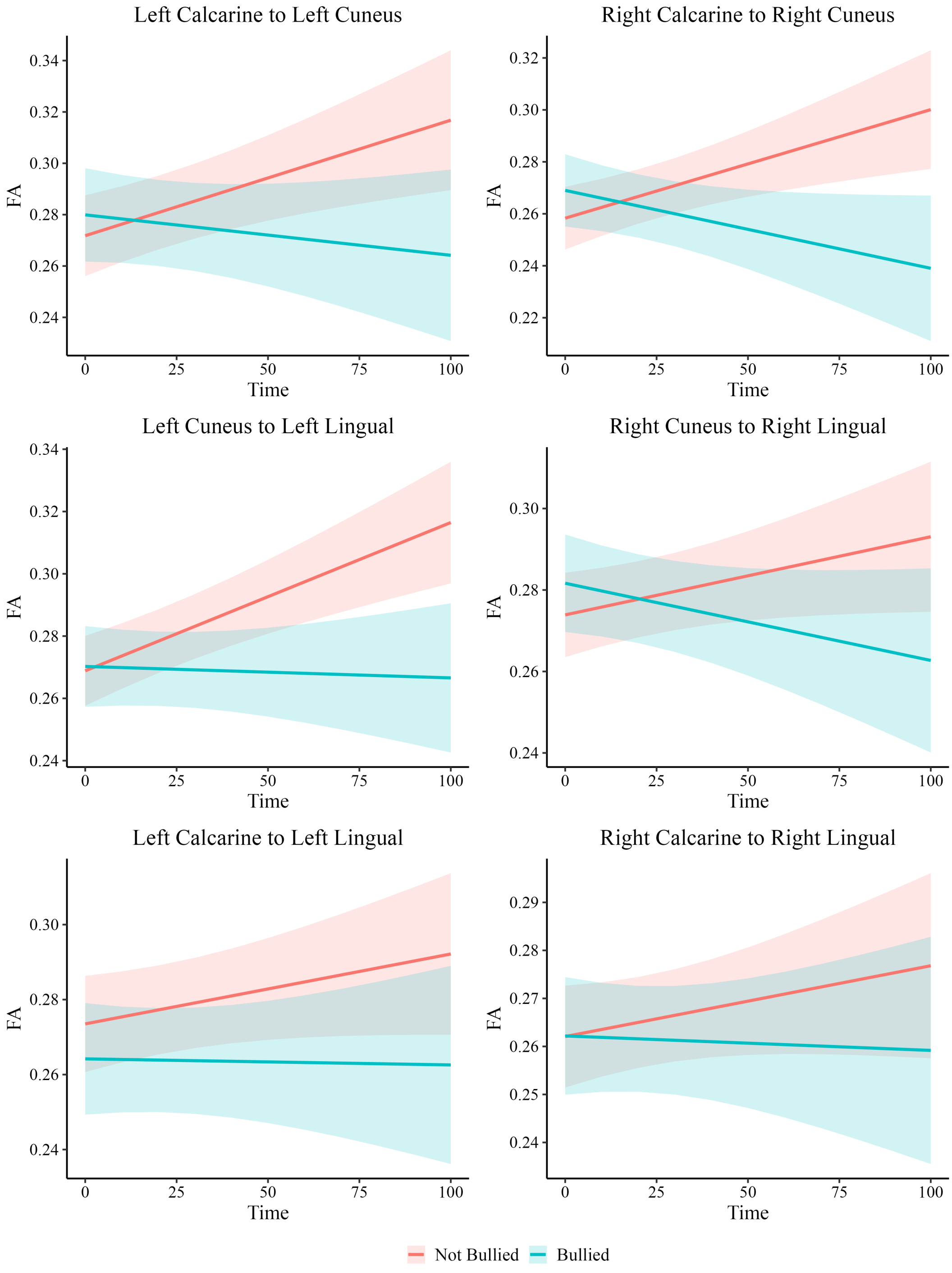

**Legend:** Predicted fractional anisotropy (FA) trajectories across time for six medial occipital white matter connections, stratified by bullying Victimisation. Time represents months since the baseline scan. Coral lines represent participants who were Not Bullied, and cyan lines represent participants who were Bullied. Shaded areas indicate 95% confidence intervals. Estimates are adjusted for sex, baseline age, total stress, and family stress.

**eFigure 2:** Medial Occipital White Matter FA Trajectories by Bullying Victimisation and Psychotic-Like Experiences.

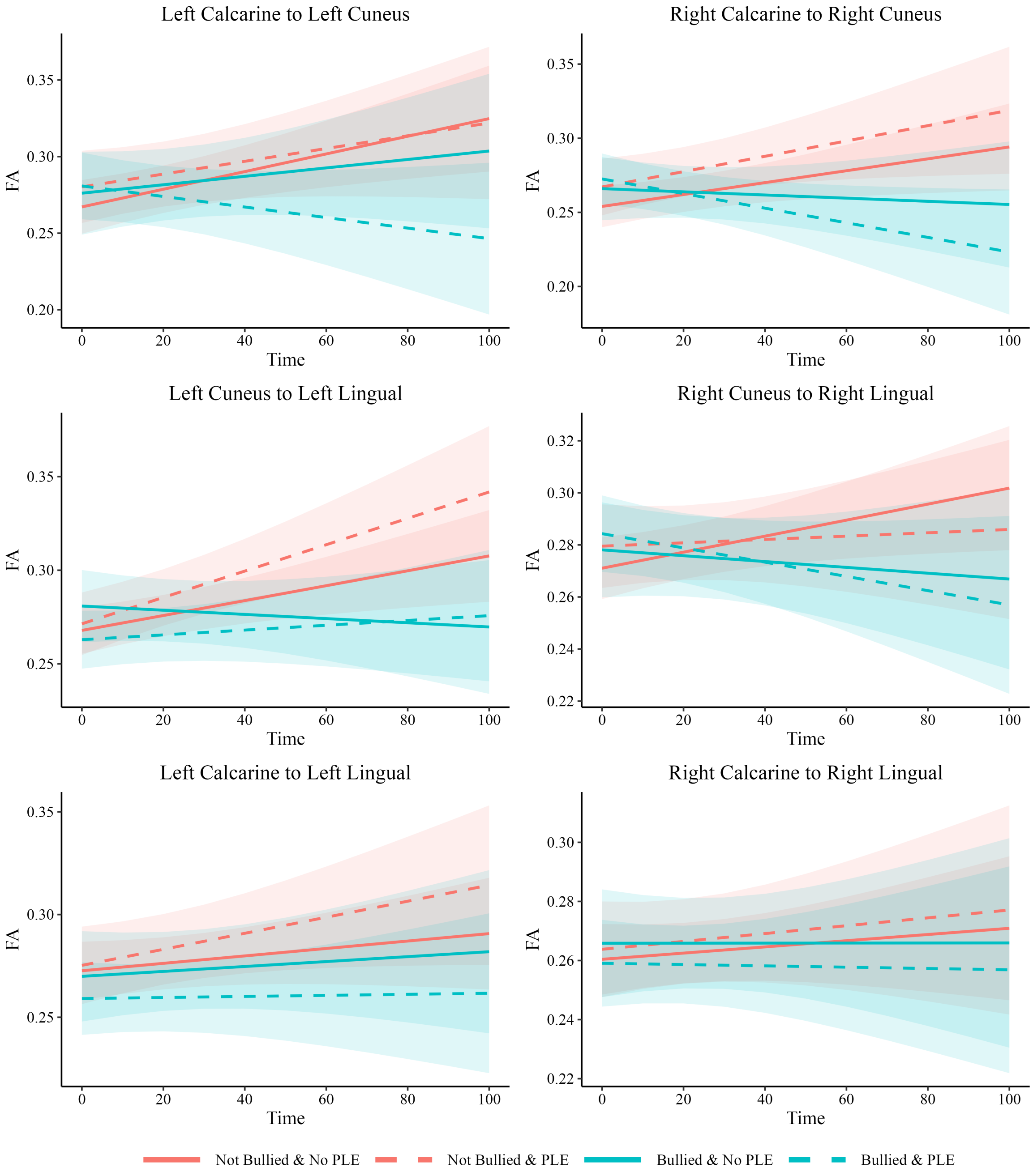

**Legend:** Predicted fractional anisotropy (FA) trajectories across time for six medial occipital white matter connections, stratified by bullying Victimisation and psychotic-like experience (PLE) status. Time represents months since the baseline scan. Coral lines represent participants who were Not Bullied, and cyan lines represent participants who were Bullied. Solid lines indicate No PLE, whereas dashed lines indicate PLE. Shaded areas represent 95% confidence intervals. Estimates are adjusted for sex, baseline age, total stress, and family stress.
